# Pollen foraging mediates exposure to dichotomous stressor syndromes in honey bees

**DOI:** 10.1101/2024.08.20.608746

**Authors:** Sydney B. Wizenberg, Sarah K. French, Laura R. Newburn, Mateus Pepinelli, Ida M. Conflitti, Mashaba Moubony, Caroline Ritchie, Aidan Jamieson, Rodney T. Richardson, Anthea Travas, Mohammed Arshad Imrit, Matthew Chihata, Heather Higo, Julia Common, Elizabeth M. Walsh, Miriam Bixby, M. Marta Guarna, Stephen F. Pernal, Shelley E. Hoover, Robert W. Currie, Pierre Giovenazzo, Ernesto Guzman-Novoa, Daniel Borges, Leonard J. Foster, Amro Zayed

## Abstract

Recent declines in the health of honey bee colonies used for crop pollination pose a considerable threat to global food security. Foraging by honey bee workers represents the primary route of exposure to a plethora of toxins and pathogens known to affect bee health, but it remains unclear how foraging preferences impact colony-level patterns of stressor exposure. Resolving this knowledge gap is crucial for enhancing the health of honey bees and the agricultural systems that rely on them for pollination. To address this, we carried out a national-scale experiment encompassing 456 Canadian honey bee colonies to first characterize pollen foraging preferences in relation to major crops, then explore how foraging behaviour influences patterns of stressor exposure. We used a metagenetic approach to quantify honey bee dietary breadth and found that bees display distinct foraging preferences that vary substantially relative to crop type and proximity, and the breadth of foraging interactions can be used to predict the abundance and diversity of stressors a colony is exposed to. Foraging on diverse plant communities was associated with increased exposure to pathogens, while the opposite was associated with increased exposure to xenobiotics. Our work provides the first large-scale empirical evidence that pollen foraging behaviour plays an influential role in determining exposure to dichotomous stressor syndromes in honey bees.

**Significance Statement:** Insect-mediated pollination is an important ecological process that is crucial for food production. Managed honey bee colonies are one of the most important insect pollinators, but their health has been under threat from a variety of stressors. Bee workers are primarily exposed to stressors while foraging and understanding how bee foraging preferences are related to exposure risk could provide pivotal information to improve management efforts. Here, we studied honey bee foraging preferences in relation to prominent Canadian crops and across a gradient of modified environments. We found that honey bees show distinct, measurable foraging preferences and that dietary diversity is a strong predictor of the stressors that colonies are exposed to.

## Introduction

The western honey bee, *Apis mellifera*, is the most frequent floral visitor of crops worldwide (1, 2), playing a vital role in the productivity of agricultural systems (3). Recent declines in the health of managed honey bee colonies, seen most prominently as an increase in annual colony losses (4), presents a considerable threat to global food security. No singular entity has emerged as a primary driver of colony losses (5, 6), but rather, poor health as a result of exposure to a multitude of stressors (7) has been suggested as a possible cause. These stressors include pathogens (8, 9), pests (10), xenobiotics (11–13), and poor-quality diets (14–16). Pin-pointing individual stressors that induce colony loss has been difficult (17,18), and attempts to model the specific mechanisms by which stressor exposure induces colony mortality have failed to produce meaningful management strategies (19, 20). Some work has suggested that pests and pathogens could play the largest role (21), but more broadly, declines in bee health have been attributed to the entangled effects of multi-stressor exposure (22), the complexities of which are only beginning to be explored (7). While these general stressors have been known for some time, it is not immediately obvious how foraging behaviour influences patterns of exposure.

Foraging bees must travel outside of the colony to collect pollen and nectar, and through this process they may be exposed to xenobiotics, pests, and pathogens. Pollen is a primary food source, providing access to important nutrients for foraging bees and the colonies they support (23–27). Colony-level foraging decisions generally occur in advance of mass pollination events (28), e.g. foraging scouts may locate ideal food sources and communicate this to the rest of the colony via a waggle dance, and are often driven by cost-benefit ratios that favor high quality resources in close proximity to the hive (29). Both the abundance and diversity of floral resources within a landscape shape these patterns of interaction; complex environments support shorter foraging distances (30, 31), but can also promote the rapid discovery and abandonment of food sources (32). Contrastingly, homogenous agroecosystems support a wider geographic foraging range (33) and a decrease in the frequency of waggle-dances (30), indicating a reduction in the discovery of new food sources and thus the richness of plant-bee interactions. Despite its obvious importance to stressor exposure, we know little about how the foraging preferences of worker bees mediate colony-level exposure to stressors. Resolving this knowledge gap is crucial for mitigating threats to honey bee health and increasing the resilience of agricultural systems that depend on honey bees for pollination.

Previous work has demonstrated that honey bees can exhibit measurable foraging preferences in relation to crop species (34–37), but this type of work has long been inhibited by the logistically constrained task of quantifying plant-pollinator interactions. Recent advancements in methodological development (38–42) have provided a new avenue for exploring plant-pollinator interactions and present an opportunity to resolve knowledge gaps about foraging behaviour and its influence on stressor exposure. Here, we set out to quantify honey bee foraging preferences in relation to prominent Canadian crops, and explore how patterns of plant interaction drive exposure to the pathogens and xenobiotics nested within anthropogenically-modified environments. Applying a metagenetic approach to characterize honey bee dietary breadth (38), we asked: do honey bees have distinct foraging preferences when presented with common Canadian crops? Does the structure of a landscape impact foraging behaviour? And finally, does foraging behaviour mediate exposure to stressors?

## Results

### Experimental overview

We used an established experimental design (13) to characterize honey bee foraging preferences, and explore how foraging behaviour mediates exposure to the stressors nested within Canadian landscapes, (Fig. 1). At the start of the beekeeping season, we randomly assigned colonies to apiaries near (i.e. in or directly adjacent) or far (>1,500 m) from the following 8 common Canadian crops: cranberry, lowbush blueberry, highbush blueberry, apple, commodity canola, hybrid seed-production canola, corn, and soybean. We sampled pollen from these colonies at two time points during the experiment (see methods) and subjected these samples to multi-locus pollen metabarcoding to identify the source and relative abundance of different plant taxa in each sample following established methods which have been demonstrated to provide results comparable to microscopic melissopalynology (38). Overall, our pollen metabarcoding libraries were deep sequenced to an average depth of 11,210,193 (± 2,298,792) barcode reads and identified over 480 plant genera, including the 5 associated with our focal crops (*Vaccinium*, *Malus*, *Brassica*, *Zea*, and *Glycine*). We additionally tested the pollen, nectar, and nurse bees from each colony for chemical residues, and the nurse bees for pathogens (7). Dietary values presented below are the multi-locus average relative abundance of reads associated with the genera for each relevant focal crop. Statistical significance was determined using a threshold of p < 0.05.

**Figure 1:**
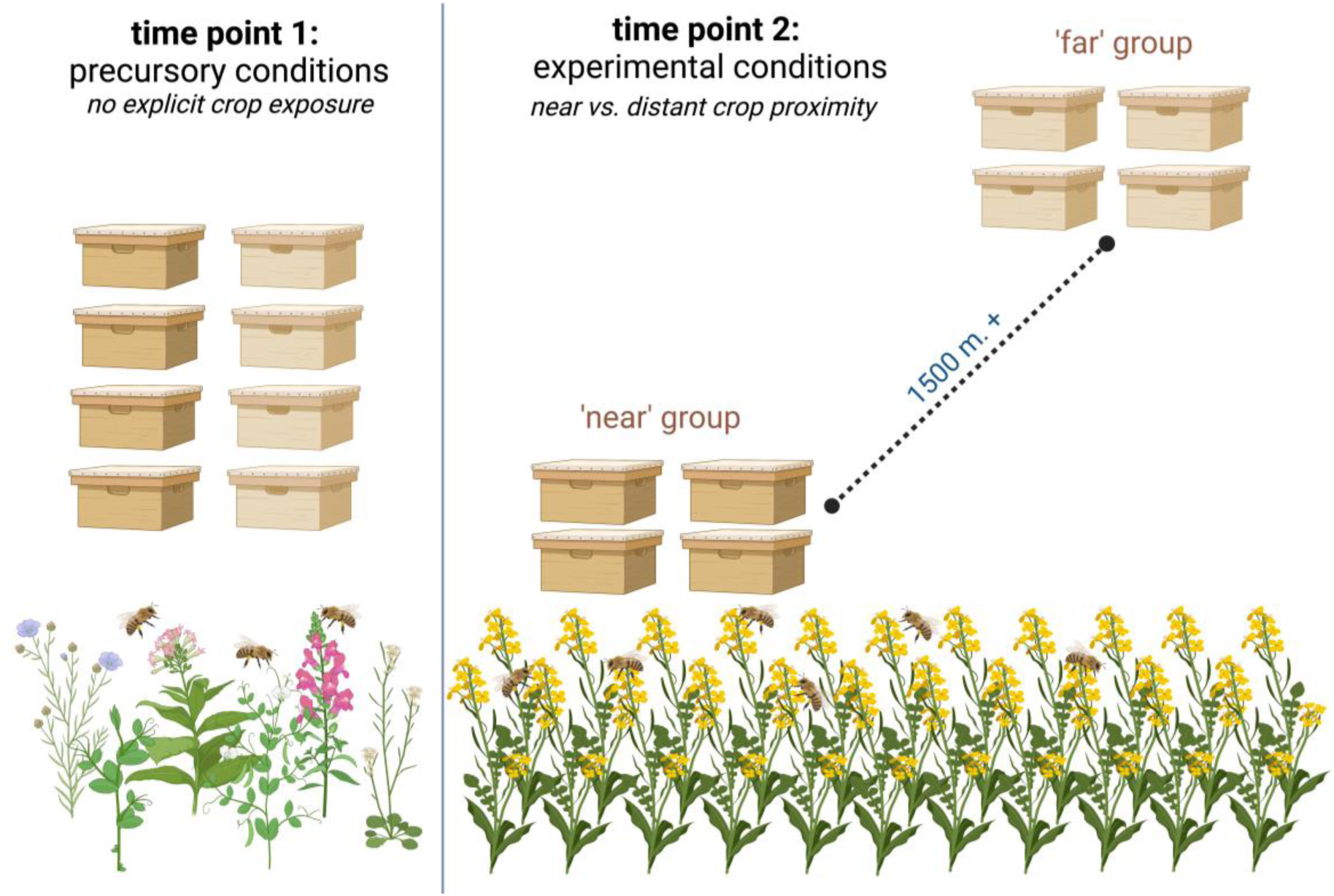
Experimental design. At the start of the season, colonies were randomly assigned to apiaries that were either ‘near’ or ‘far’ from focal crops. Colonies were sampled after randomization prior to crop exposure (time point 1) and during periods of maximum exposure (time point 2). Image created with BioRender.com.

### Honey bees display strong foraging preferences

The 456 honey bee colonies used in our experiments displayed strong foraging preferences – quantified by pollen metabarcoding – in relation to common Canadian crops. Only 4 of our 12 experiments had a “proximity-dependent” effect wherein focal crop pollen dietary abundance (the relative dietary proportion comprised of the genera associated with the focal crop being studied) differed significantly between colonies placed near and far from the focal crop (S1: Table S1, Table S2). In both of our cranberry experiments, as well as two of our blueberry experiments, we detected a statistically significant difference in the relative dietary abundance of *Vaccinium* pollen between our near and far groups. When a honey bee colony was placed directly adjacent to cranberry crops, cranberry pollen comprised an average of 28.5% (± 21.8% SD) of their diet, and when placed far away from cranberry crops, we detected no cranberry pollen in their diets. This indicates that cranberry pollen is a floral resource honey bees may forage on only when it’s convenient, e.g., spatially close and abundant (Fig. 2). Blueberry crops were not a highly accessed floral resource over-all. Even when colonies were placed directly adjacent to the crop, blueberry pollen only comprised an average of 3.2% (± 3.6) of their diet. This indicates that blueberry pollen may not be a favorable dietary component, but honey bees will forage on it to some degree if its easily accessible (Fig. 2). Across all canola experiments, distance to the focal crop (near or far) had no discernable impact on foraging preferences, and canola comprised a substantial (47.9% ± 21.6) portion of each colonies’ diet regardless of the relative proximity of the crop (Fig. 2). Similarly, apple foraging behaviour did not differ between near and far groups, but only comprised a small dietary proportion (5.9% ± 5.8), indicating that it was not as highly favored as other crops like canola. Both corn and soybean pollen were rarely, if ever, detected in pollen samples from these experiments (Fig. 2). Notably, our pollen analysis does not indicate the frequency of nectar feeding, and thus we cannot make any conclusions about this type of foraging interaction.

**Figure 2:**
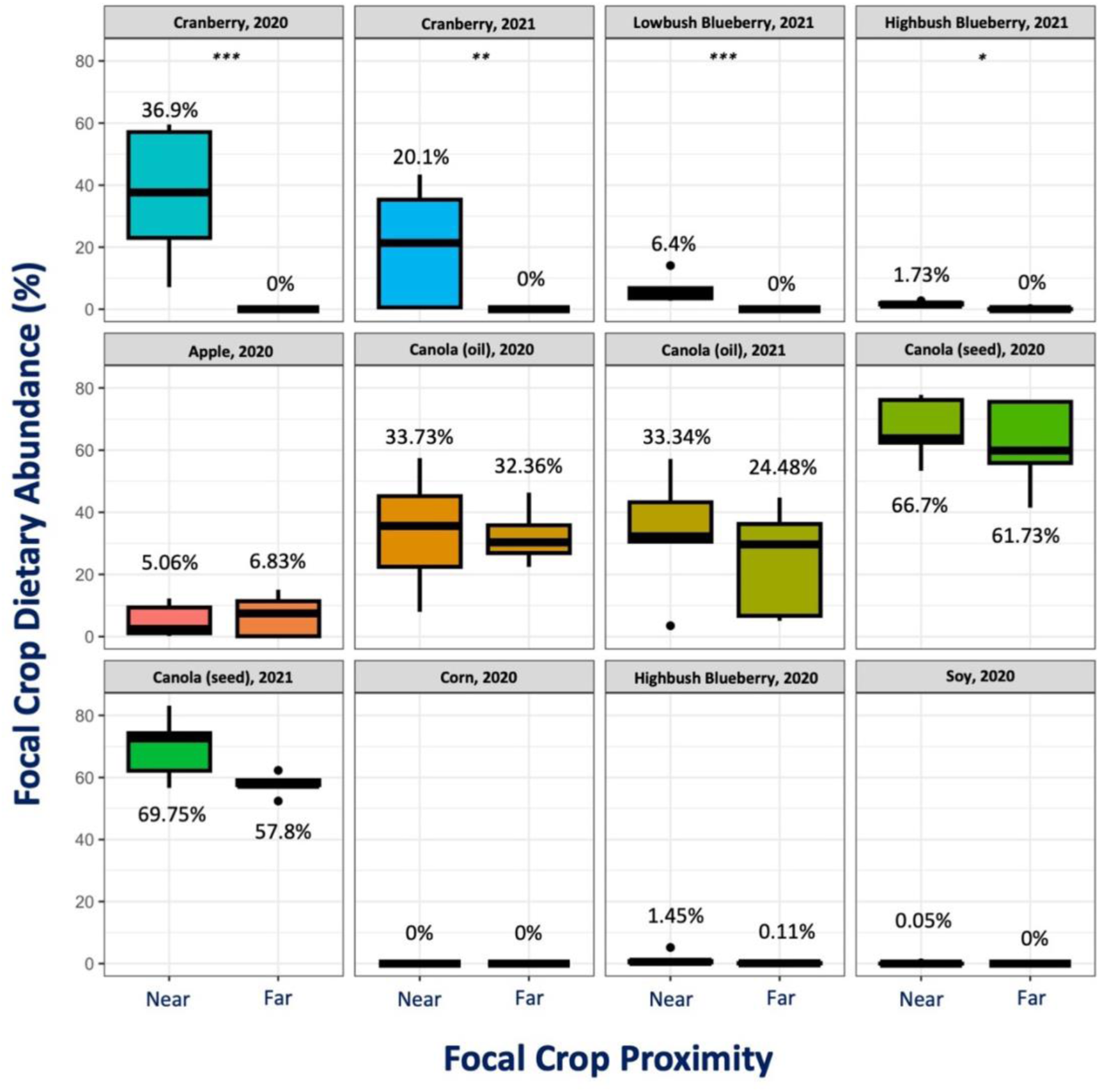
Bees display distinct foraging patterns. Boxplots show differences in focal crop pollen abundance between near and far sites. Focal crop dietary abundance was estimated as the multi-locus average relative proportion of each pollen sample comprised of the genera associated with the crop of interest. Asterisks indicate statistically significant differences (*, p < 0,05; **, p < 0.01; ***, p < 0.001), calculated using Kruskal-Wallis non-parametric analysis.

### Landscape composition predicts dietary diversity

We next explored how the relative abundance of land cover types at each site influenced dietary diversity – a quantification of the variety and abundance of plant genera within each mixed pollen sample (S1: Fig. S1). We estimated dietary diversity via Shannon Weaver’s index of species diversity and used it as an indicator of the breadth of foraging interactions; higher diversity values were strongly correlated with a low predominant dietary component and high number of unique plant genera (S1: Fig. S1). Agricultural land cover, urban land cover, and grassland cover were all significantly associated with dietary diversity (Table 1). Agricultural land was negatively correlated with dietary diversity (Fig. 3a), while urban land was positively associated with dietary diversity (Fig. 3b). Grassland was negatively associated with dietary diversity, however, this relationship was driven by a low number of observations in a single province (Fig. 3c). Forest land cover displayed no detectable association with dietary diversity (Fig. 3d).

**Figure 3:**
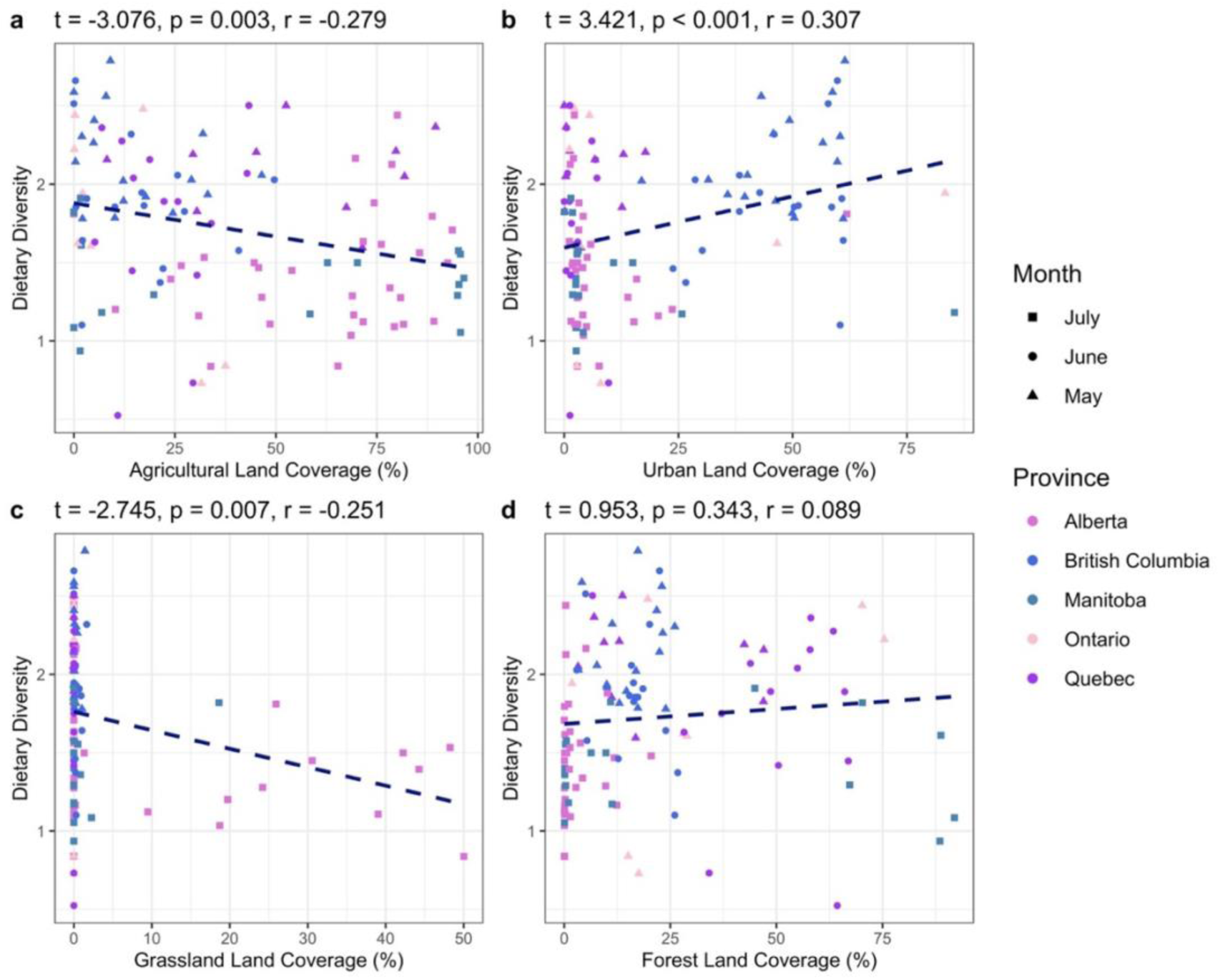
Landscape composition is significantly associated (p < 0.05) with dietary diversity. Pearson’s product-moment correlation tests assessed raw association between landscape types and dietary diversity. Land coverage (%) values indicate the relative abundance of a land-type within a 1500 m. radius of the study site. Agricultural land cover was negatively associated with dietary diversity, urban land cover was positively associated with dietary diversity, and grassland cover was negatively associated with dietary diversity (note: grassland observations were limited to a single province); t is the test statistic, r is the Pearson correlation coefficient, and p-value indicates the significance of association. Dashed lines indicate the direction of association.

**Table 1:**
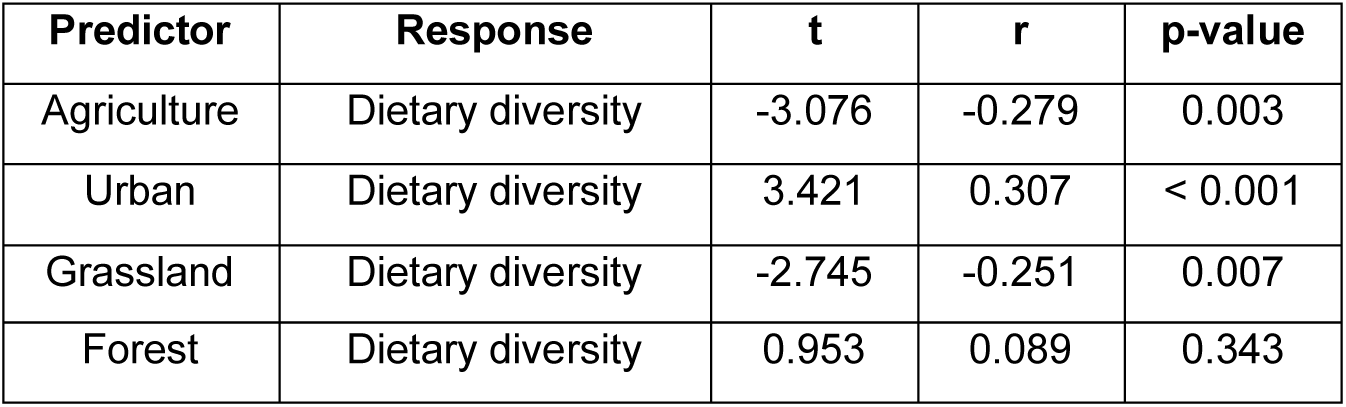
Landscape composition is significantly associated with dietary diversity (p < 0.05), analyzed via Pearson’s product-moment correlation tests. Output of product-moment correlation tests exploring association between landscape composition and dietary diversity. Agricultural land cover was negatively associated with dietary diversity, urban land cover was positively associated with dietary diversity, and grassland cover was negatively associated with dietary diversity (note: grassland observations were limited to a single province); t is the test statistic, r is the Pearson correlation coefficient, and p-value indicates the significance of the relationship.

### Dietary diversity predicts stressor exposure

Our stressor dataset was heavily skewed by low or no observations, and thus we selected a subset of variables with sufficient observations (> 25 detections) to undergo mixed-effect logistic modelling, using experiment (time, location) as our random effect. Of the 18 stressors deemed suitable for logistic analysis, 8 were significantly associated with dietary diversity, assessed via log-odds estimates and p-values (Table 2). Log-odds ratios indicated that as dietary diversity decreased, we were significantly more likely to detect 4 xenobiotics: coumaphos, picoxystrobin, clothianidin, and thiamethoxam (Fig. 4; Table 2). In contrast, as dietary diversity increased, we were significantly more likely to detect *Nosema* spores, European Foulbrood, as well as two xenobiotics: flupyradifurone, and pyrimethanil. (Fig. 4; Table 2). Five stressor variables had both a sufficient number of observations and variation in those observations to undergo general linear mixed-effect modelling: *Nosema* spore abundance, Varroa mite abundance, summed total viral loads, thiamethoxam LD_50_ risk quotients (RQs), and clothianidin RQs. LD_50_ risk quotients quantify the relative risk of adverse effects of exposure given the observed concentration, estimated dietary intake, and the known lethal dose (see methods for a description of these calculations). Due to the non-normal distribution of stressor observations, we opted to use quasi-poisson mixed-effects models to better account for the high number of low or no detections. Both neonicotinoid LD_50_ RQ’s were significantly negatively associated with dietary diversity (Table 3); the highest observations of both thiamethoxam and clothianidin were seen in colonies with low dietary diversity (Fig 5a,b). In contrast, total viral loads were significantly positively associated with dietary diversity (Table 3); colonies with high dietary diversity tended to have the greatest total pathogen loads (Fig 5c). To determine if the association between dietary diversity and quantitative stressor abundance was confounded by the influence of landscape composition, we ran the same quasi-poisson mixed-models, using relevant land cover parameters as the model input. Landscape composition was a weak predictor of stressor abundance (Fig. 5d,e,f), suggesting that our dietary diversity models are not substantially confounded by the known influence of landscape composition.

**Figure 4:**
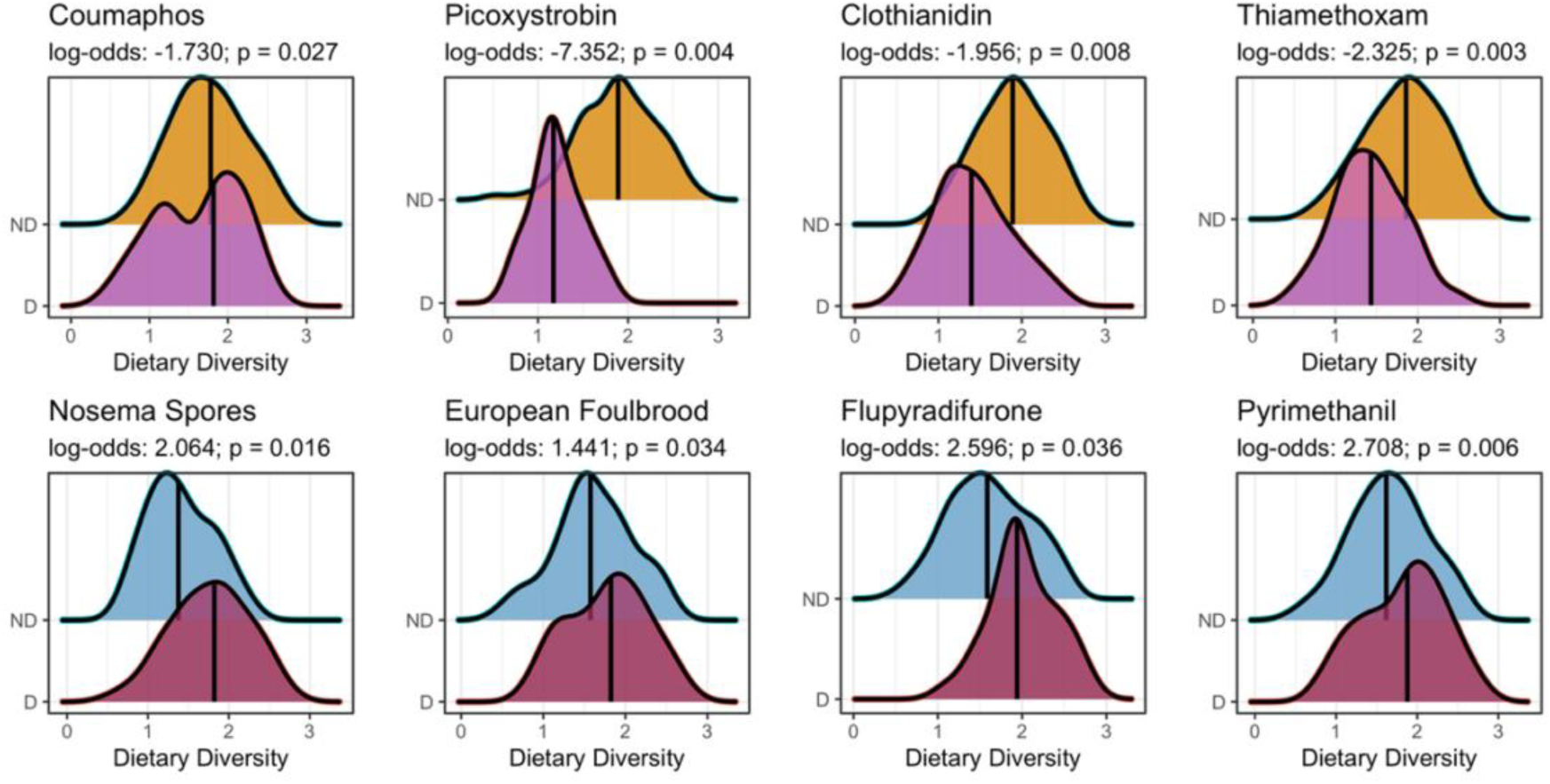
Dietary diversity is a strong predictor of stressor exposure. Ridge plots display significant association (p < 0.05) between dietary diversity and the detection outcome of 8 stressors, analyzed via mixed-effect binomial logistic regressions. Log-odds ratios indicate a change in the probability of detection as dietary diversity increases; negative values indicate a reduction in the likelihood of exposure, positive values indicate an increase in the likelihood of exposure.

**Figure 5:**
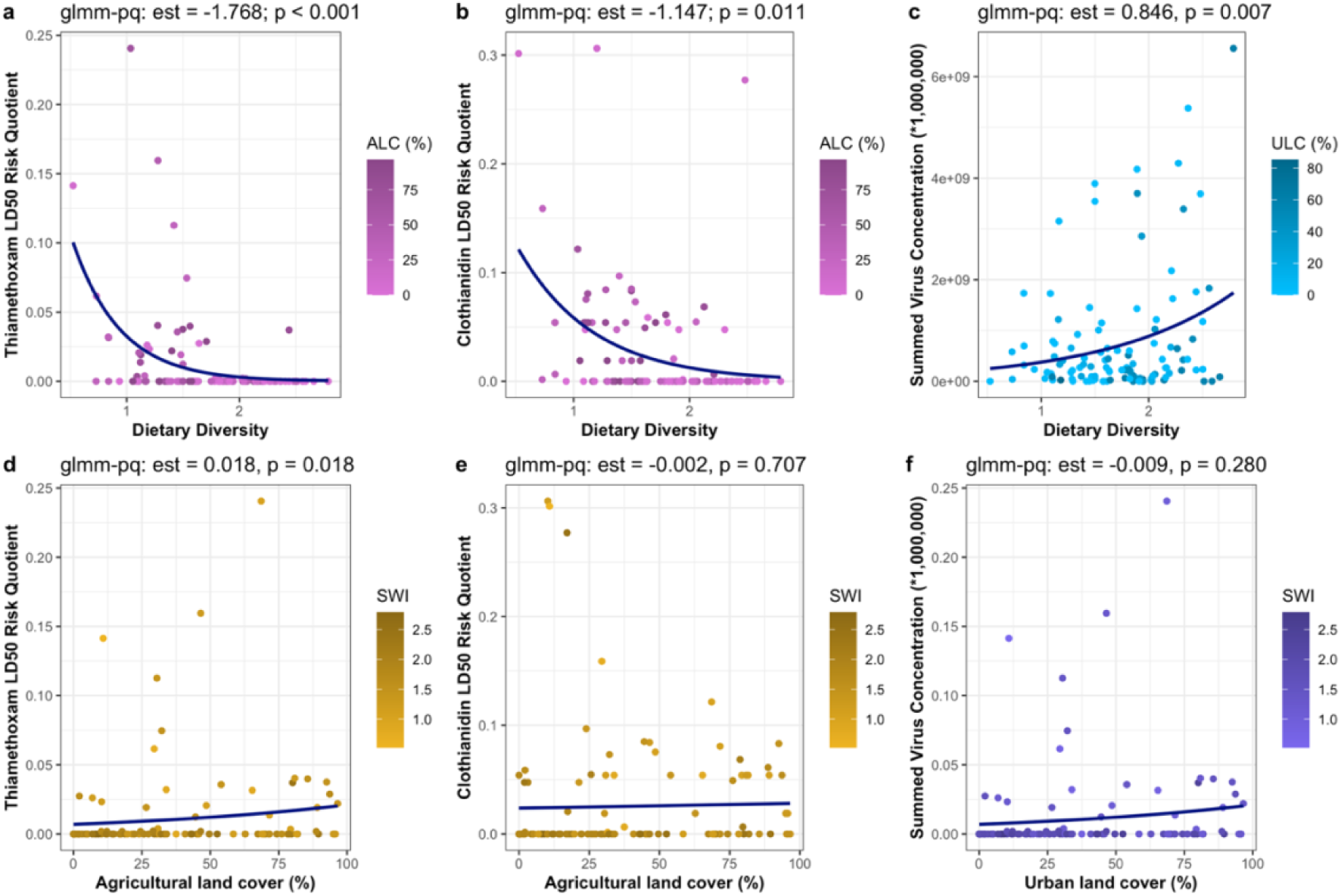
Dietary diversity is a significant predictor of stressor abundance. Quasi-poisson mixed effect models (glmm-pq) tested for association between the response (stressor abundance), and the predictor (dietary diversity, or landscape composition), controlling for experiment (location, time) as a random effect. Estimates are quasi-poisson log-odds coefficients, p-values indicate the statistical significance of the relationship. Dietary diversity (a,b,c) was a significant predictor (p < 0.05) of stressor exposure, while relevant landscape parameters (d,e,f) were weak predictors of stressor exposure. ALC (agricultural land coverage) and ULC (urban land coverage) are landscape gradients values that indicate the relative abundance of a land-type within a 1500 m. radius of the study site. SWI (Shannon’s Index) are dietary gradients that indicate the relative diversity of a pollen sample.

**Table 2:**
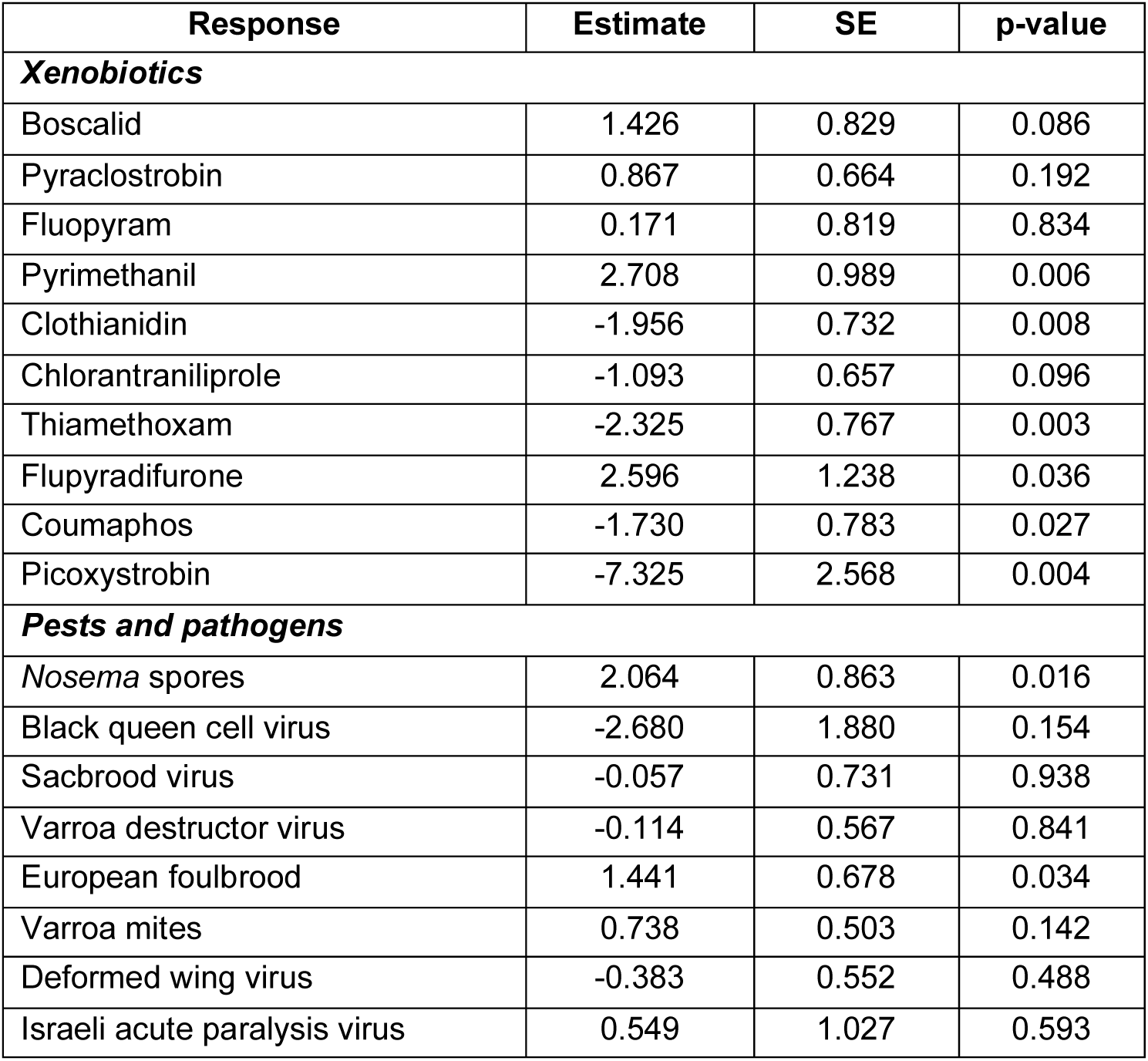
Dietary diversity is associated with the detection of 8 stressors of interest, analyzed via mixed-effect binomial logistic regressions. Output of mixed-effect binomial logistic regressions testing for association between dietary diversity and the presence or absence outcome of a stressor response. Estimate is the log-odds of detection as dietary diversity increases, SE is the standard error of that estimate, p-values indicate the statistical significance of the relationship.

**Table 3:** Dietary diversity is associated with quantitative stressor levels, analyzed via quasi-poisson mixed-effect linear regressions. Output of quasi-poisson mixed-effect linear regressions testing for association between the response (stressor abundance), and the predictor (dietary diversity, or landscape composition), controlling for experiment (location, time) as a random effect. Estimates are quasi-poisson log-odds coefficients, p-values indicate the statistical significance of the relationship.

| Predictor | Response | Estimate | SE | t-value | p-value |
| --- | --- | --- | --- | --- | --- |
| ***Diet predicting stressor abundance*** |  |  |  |  |  |
| Dietary diversity | *Nosema* spores | 0.321 | 0.207 | 1.549 | 0.124 |
| Dietary diversity | Mites | 0.266 | 0.342 | 0.778 | 0.438 |
| Dietary diversity | Viral load | 0.846 | 0.308 | 2.751 | 0.007 |
| Dietary diversity | Thiamethoxam | -1.768 | 0.404 | -4.377 | < 0.001 |
| Dietary diversity | Clothianidin | -1.147 | 0.442 | -2.593 | 0.011 |
| ***Landscape predicting stressor abundance*** |  |  |  |  |  |
| Urban land | Viral load | -0.009 | 0.008 | -1.086 | 0.280 |
| Agricultural land | Thiamethoxam | 0.018 | 0.008 | 2.413 | 0.018 |
| Agricultural land | Clothianidin | -0.002 | 0.007 | -0.376 | 0.707 |

## Discussion

The honey bee colonies used in our experiments exhibited distinct foraging preferences; canola was a highly sought-after floral resource, while cranberry foraging appeared to be driven by convenience, and other crops (blueberry, corn, soybean) were clearly not a dominant dietary source of pollen. There was a strong preference for both canola production systems in our experiments, especially the seed production variety; even colonies placed > 1500 m from flowering canola had diets dominated by *Brassica* pollen. This is in sharp contrast to the lack of interest in other crop genera - even when placed directly adjacent to blueberry plants during anthesis, pollen stored by honeybees rarely contained any *Vaccinium* pollen, implying that they may be actively foraging outside of their expected range to access preferred floral resources. One crop in particular was only notably foraged on by ‘near’ groups: cranberry, indicating a moderate preference that is potentially driven by convenience. The strong foraging preference for canola seen in our experiments coincides with that seen by Rollin et al. (36), as does the indifference to blueberry species seen by Girard et al. (37). Interestingly, our experiment documented no pollen-foraging of corn by honey bees, in direct contrast to the results seen by Danner et al. (35), but coinciding with the results seen by Tsvetkov et al. (12).

To some degree, foraging patterns may be explained by pollen dispersal mechanisms - corn is anemophilous, relying on wind as the predominant vector for pollen dispersal (43, 44). Anemophilous (wind-dispersed) species are unlikely to comprise a substantial proportion of any honey bees’ diet due to their small size and lack of pollenkitt, an adhesive compound found on the outer exine layer that aids in the dissemination of entomophilous (insect-dispersed) pollen (45, 46). Soybean may not rely on anemophily for dispersal of pollen, but some work has suggested it predominantly self-pollinates (47), and thus may not invest resources into the attraction of insect pollinators. Dispersal mechanisms provide a simple explanation for some foraging patterns but fail to account for the variable behaviour seen in our cranberry and blueberry experiments. This lack of interest in blueberry pollen has been seen previously (37), wherein the authors propose that it may be the result of a poor nutrient profile. More broadly, blueberry is thought to require buzz pollination, i.e. sonication of flowers to promote anther dehiscence (48), which honey bees are not known to do (49, 50). Thus, low dietary abundance could be a result of insufficient sonication abilities, coupled with low over-all pollen production rates. Alternatively, if nutritional demands really are driving foraging patterns, the consistent treatment of canola (*Brassica*) as a high-value and highly accessed floral resource could imply that it has special dietary value. Experimental work has suggested that diets comprised of *Brassica* pollen can reduce early mortality of worker bees (51), though its protein:lipid ratio is relatively low (52), contradicting the hypothesis that preferences are driven by protein content (53). Evidently, more work is needed to understand the specific mechanisms that determine foraging value and the related preference for canola pollen.

Across all 12 experiments and 114 sites, our work has provided robust evidence that within Canadian landscapes, diets of lower diversity are associated with greater exposure to xenobiotics (Fig. 4), specifically the neonicotinoids thiamethoxam and clothianidin (Fig. 5a,b), while diets of higher diversity are associated with greater exposure to pathogens (Fig. 4; Fig. 5c). These findings demonstrate that pollen foraging behaviour and related patterns of plant interaction drive exposure to dichotomous stressor syndromes, characterized by diets of low diversity and increased xenobiotic exposure, or diets of high diversity and increased pathogen exposure. Landscape composition is evidently influential in determining these patterns of interaction (Fig. 3), and while the relationship between landscape composition and dietary diversity was significant, the associated correlation coefficients were weak (Table 1). Further, landscape composition alone was a weak predictor of stressor abundance (Fig. 5d,e,f), demonstrating the key role that foraging preferences play in mediating exposure. Pollen foraging preferences providing linkage between the structure of a landscape and exposure to the stressors nested within that landscapes is intuitive, and yet, our work is the first to empirically document this on a large scale.

Our findings suggest that patterns of stressor exposure occur on a dichotomous scale, and the degree to which any given colony experiences one of these contrasting stressor syndromes is mediated by patterns of plant interaction. Foraging on singular pollen sources within agricultural contexts led to increased xenobiotic, and specifically neonicotinoid, exposure – highlighting the risks associated with crop pollination and its impact on colony health. Though our work is one of the first to provide direct evidence of this linkage, the risks associated with crop pollination have been known for some time. Crop monocultures are routinely treated with xenobiotics (54–56), such as the neonicotinoid thiamethoxam (Fig. 4; Fig. 5a), which is known to impair flight ability (57), alter motor function (58), and cause oxidative damage (59). Similar deleterious effects have been documented after exposure to clothianidin (60–62), the detection of which was again associated with low dietary diversity (Fig. 4; Fig. 5b). Though it remains hotly debated how detrimental neonicotinoid exposure may be to honey bee health, recent work has demonstrated the degree of impact may be genotype dependent (63), suggesting that some colonies may be more susceptible than others. While crop pollination remains necessary, developing strategies to reduce colony reliance on singular pollen sources could present a pathway to reduce the degree of xenobiotic exposure. Agroecosystems populated by crop monocultures provide little diversity of pollen sources to foraging bees, and thus could be the root cause of this linkage. It remains unclear if low dietary diversity is associated with nutritional stress, but if this hypothesis holds true, honey bee colonies foraging on crop pollen may be more likely to experience the antagonistic effects of multi-stressor exposure via the compounded exposure to xenobiotics and a poor-quality diet.

Increasing the diversity of floral resources available to foraging bees may reduce singular reliance on crop pollen, and any associated crop-specific xenobiotics, but this strategy comes with its own set of risks. Our work suggests that diets with a diverse mix of pollen sources are associated with increased pathogen exposure (Fig. 4; Fig. 5c), representing an alternative but potentially equally stressful scenario for managed honey bee colonies. This foraging behaviour was promoted by urban land cover (Fig. 3b), coinciding with previous findings that suggest that cities promote high foraging diversity, likely as a result of high beta plant diversity and fine-grain heterogeneity of floral resources (64). Interestingly, other work has provided evidence that cities, heavily comprised of impermeable surfaces (65–67), can concentrate invertebrates into small densely-populated green spaces, facilitating the development of pathogen transmission hubs (68). This is likely attributable to the overlap in resource access patterns when the availability of floral resources is limited; interspecific interactions are a primary source of microorganism transmission (69–72). Disease spill over from honey bees has led to the spread of pathogens among other bee taxa (73), primarily via overlap in plant interactions (69, 74), and while honey bees may have been the catalyst for the initial introduction of many microorganisms, transmission is multilateral. Our work provides linkage between these findings, suggesting that cities promoting diverse foraging behaviour and increased pathogen exposure are an intertwined phenomenon. It is possible that pollinator gardens and other pockets of biodiversity within urban developments could act as poisoned oases (75), providing diverse diets at the expense of exposure to pathogens.

Here, we present the first large-scale evidence that pollen foraging preferences in modified environments mediate exposure to dichotomous stressor syndromes. Colonies foraging at agricultural sites on a small number of pollen sources experience a greater risk of xenobiotic exposure, while colonies foraging at urban sites on diverse pollen sources experience a greater risk of pathogen exposure. The dichotomous nature of stressor exposure found in our 456 honey bee colonies suggests that modified landscapes, regardless of the nature of those modifications, present a broad risk to honey bee health. This implies that despite our best efforts, no anthropogenically modified environment is wholly “safe” for honey bee colonies, and supporting pollinators may require a radical change in our urban and agricultural land management strategies. What remains largely unknown is how dietary diversity (e.g., nutrition) impacts colony health - integrated with our findings here, this key question could provide crucial information to improve our efforts to combat colony loss and maintain healthy, robust colonies to pollinate our crops and increase the resilience of our food production systems.

## Materials and Methods

### Experimental design

We used an established experimental design (7,13) to explore how crop proximity impacted the frequency of plant interaction in 8 Canadian crops: cranberry (*Vaccinium*), highbush blueberry (*Vaccinium*), lowbush blueberry (*Vaccinium*), commodity canola (*Brassica*), hybrid seed-production canola (*Brassica*), apple (*Malus*), soybean (*Glycine*), and corn (*Zea*). Crops of particular interest (both canola production systems, highbush blueberry, cranberry) were replicated across a second year, totaling 12 spatially and/or temporally separate experiments. Each experiment followed the same general design (Fig. 1); at the beginning of the pollination period for each crop, we placed 40 honey bee colonies at a neutral apiary and sampled under ‘t1’ precursory conditions (described below) to confirm that no experimental group differed substantially in their foraging behaviour or stressor load preceding the start of each experiment. We selected colonies of equal size and visually inspected them to ensure that they were healthy with no obvious signs of disease prior to the start of the experiments. We then randomly assigned each of the colonies to one of 10 experimental sites. Five of those sites were in or directly adjacent to the focal crop of interest (‘near’ crop proximity), and the remaining 5 were placed a minimum of 1500 m away from the focal crop (‘far’ crop proximity), with all sites being at least 1500 m away from another experimental site. After moving the colonies to sites, we performed ‘t2’ sampling (experimental conditions) 2-4 weeks after placement, coinciding with the timing of peak blooming for the relevant focal crop. We do not provide exact GPS coordinates for apiaries to protect the identity of the beekeepers and farmers involved. The approximate location (municipality, province) of each site is provided in a supplemental file (S2).

### Sampling

At each time point (t1 precursory conditions and t2 experimental conditions), we sampled several colony matrices to quantify biotic and abiotic stressors and the composition of stored pollen (bee bread). We sampled 4 g of bee bread (or approximately 100 hive cells) from each of 4 colonies at a site by harvesting freshly processed bee bread, identifiable by the colour and light degree of compaction. We then pooled all 4 samples, and divided the resulting 16 g sample into two portions: 10 g to undergo pollen metabarcoding (described below), and 6 g to undergo xenobiotic analysis (described below). We next sampled nectar from each colony at each site; we collected 3 mL from the comb, using a 1-cc syringe with the needle removed, and transferred it to a 5 mL centrifuge tube. Nectar was used for xenobiotic analysis (described below). We then sampled nurse bees for pathogen and xenobiotic analysis. We randomly sampled 200 bees from each site (∼ 50 per colony), which were placed on dry ice and stored in a -80 °C freezer. We quantified *Varroa destructor* mite abundance using an alcohol wash method (77) and standardized varroa mite counts for each colony to represent the number of mites per 100 bees, which was then averaged across the 4 colonies at each site and time point.

### Pathogen quantification

We quantified a number of pathogens in the collected samples of hive bees using established methods (76, 78) performed by the National Bee Diagnostic Centre (Beaverlodge, Alberta). This included qPCR quantification of acute bee paralysis virus, black queen cell virus, chronic bee paralysis virus, deformed wing virus, Israeli acute paralysis virus, Lake Sinai virus, Kashmir bee virus, sacbrood virus, and *Varroa destructor* virus-1. The causative agent of European foulbrood (*Melissococcus plutonius*) was detected using qPCR20 with a 7500 Fast Real-Time PCR System (Applied Biosystems, Foster City, USA). *Nosema* spores were quantified using microscopy (78). Each sample was first homogenized in 12 mL of GITC (guanidinium thiocyanate) buffer. We then isolated RNA using a 200 μL aliquot and the NucleoSpin®RNA kit (Macherey-Nagel Gmbh & Co. KG, Düren, Germany), and used 800 ng to synthesize cDNA at 46°C for 20 min using the iScript cDNA synthesis kit (Bio-Rad laboratories, Hercules, USA). The resulting 20 μL of dDNA was diluted with 60 μL of nuclease-free sterile water, and 3 μL were used for qPCR analysis. Our quantitative PCR analysis of each sample to determine the presence and abundance of pathogens of interest used SSoAdvanced™ Universal SYBR® Green Supermix (Bio-Rad Laboratories, Hercules, USA) and the previously published primers described by Borba et al. (78). Triplicate assay amplifications used approximately 30 ng of cDNA, and a CFX384 Touch™ Real-

Time Detection System (Bio-Rad Laboratories, Hercules, USA). Standard curves were generated from plasmids that contained the target sequences with diluted copy numbers ranging from 10^2^ to 10^7^. PCR program specifications were as follow: initial denaturation for 3 min at 95°C, followed by 10 s at 95°C and 20 s at 60 °C for 40 cycles. We confirmed the accuracy of these approaches via melt-curve analysis (65–95°C with increments of 0.5°C at 2 s/step), real-time qPCR data was analyzed with the CFX Manager™ Software (Bio-Rad Laboratories, Hercules, USA).

#### Xenobiotic quantification

Multiresidue quantification of 239 xenobiotics was carried out on samples of pollen and nectar following standard methods (79, 80) by the Agriculture and Food Laboratory (Guelph, Ontario), using a modified QuEChERS (quick, easy, cheap, effective, rugged, and safe) extraction. For each sample that underwent analysis, a two-gram sub-sample was extracted into 1% acetic acid (in acetonitrile, anhydrous sodium acetate, and magnesium sulfate), and the resulting supernatant was then evaporated and diluted in methanol with ammonium acetate (0.1 M). Extracts were analyzed via LC/ESI-MS/MS (liquid chromatography and electrospray ionization tandem mass spectrometry) using an established approach (79). To compare the relative risk of xenobiotic exposure, we calculated the risk quotient (RQ) of each chemical (81, 82) based on its relative abundance and LD_50_ (median lethal dose that kills 50% of bees in test cages), as well as the approximate weight of pollen and nectar that an in-hive honey bee consumes per day, where:

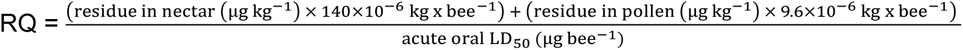

An LD_50_ was used if it was derived using workers from any subspecies of *Apis mellifera*, followed the Organisation for Economic Co-operation and Development guidelines for acute oral or diet toxicity tests on honey bees, and provided per bee or per body weight dosages. If tests using a high purity of the active ingredient were unavailable, studies on formulations or a combination of xenobiotics were used. Xenobiotic concentrations that fell below the limits of quantification or detection were assigned a concentration equal to that limit.

#### Pollen metabarcoding

We followed an established protocol for pollen metabarcoding (38). Briefly, we extracted DNA from pollen samples using the NucleoMag DNA Food Kit (Macherey-Nagel, Düren, Germany) by combining 10 g of bee bread with 20 mL of lysis buffer: 70% autoclaved filtered water (Millipore Sigma, Burlington MA, USA), 20% 10x STE (100 mM NaCl, 10 mM Tris, 25 mM EDTA), and 10% diluted SDS (10% sodium dodecyl sulfate). We then sealed each conical tube, inverted them 10 times, then homogenized the suspension by shaking samples for 10 minutes in an orbital shaker (G25 Incubator Shaker, New Brunswick Scientific, Edison, NJ, USA) at 25°C and 375 rpm. Immediately after removing the samples from the orbital shaker, we pipetted 3 mL of the homogenized sample into a 7 mL cylindrical tube containing 10 small (1.4 mm) and 2 large (2.8 mm) ceramic beads and bead beat it (Bead Mill 24, Fisherbrand, Ottawa, Ont., Canada) for 4 x 30 s cycles, at a speed of 6 m/s. We then transferred 550 uL of the homogenized suspension to a 1.5 mL Eppendorf tube. We warmed CF lysis buffer for 10 min in a 65°C water bath then added 550 uL of the warmed buffer and 10 uL of Proteinase K (from the NucleoMag DNA Food kit) to the homogenized sample and vortexed (Mini Vortex Mixer, VWR, Mississauga, Ont., Canada) the sealed tube for 30 s. We then incubated the sample at 65 °C for 30 min in a block heater (Isotemp 145D, 250V, Fisherbrand, Ottawa, Ont., Canada), inverting every 10 min. After incubation, we added 20 uL of RNase A (New England Biolabs, Ipswich, MA, USA) and allowed the sample to incubate at room temperature (20°C) for an additional 30 min. After incubation, we centrifuged the sample for 20 min at 14,000 rpm (Centrifuge 5810 R, 15 amps, Eppendorf, Hamburg, Germany), transferred 400 uL of the upper liquid layer to the binding plate, added 25 uL of NucleoMag B-Beads and 600 uL of binding buffer CB (both from the NucleoMag DNA Food kit) then ran an extraction program on the KingFisher Flex extraction robot (ThermoScientific, Waltham, MA, USA). Each of the 5 deep well plates used to complete the extraction program contained either 600 uL of CMW buffer (wash 1), 600 uL of CQW buffer (wash 2.1), 600 uL of 80% EtOH (wash 2.2), or 100 uL of buffer CE (elution). After the extraction program was complete, we transferred 80 uL of the eluted sample to a fresh 1.5 mL Eppendorf tube and froze it at -20 °C until we began DNA amplification.

We carried out three PCR programs – first amplifying the DNA barcode locus of interest, then extending the length of the amplified sequence, and finally indexing the samples with unique combinations of forward and revers primers. We used validated primers (38) to amplify loci from two barcoding regions, ITS2 and rbcL (Table 4). We used 96 well plates containing 84 pollen samples, 6 negative controls, and 6 positive controls (Banana, *Musa* sp.). Each reaction included 11 uL of water, 12.5 uL of 2x Taq Pol Mix (New England Biolabs, Ipswich, MA, USA), 0.5 uL of each relevant forward and reverse primer (1 µM), and 0.5 uL of sample DNA (∼ 36.2 ng) into each well. PCR cycling conditions were (Eppendorf Mastercycler, Ep Gradient, Hamberg, Germany): initial denaturation (94°C, 10 min, 1 cycle), followed by 40 cycles of denaturation (94°C, 30 s), annealing (54°C, 40 s), and extension (72°C, 1 min), then a final extension cycle (72 °C, 10 min). The product of this first PCR (PCR1) was used as the template for a second PCR reaction (PCR2; Table 4) and the same chemistry as described above (with 0.5 uL of PCR1 product instead of sample DNA). Cycling conditions were the same for PCR1 and PCR2, with the exception of a higher annealing temperature (56°C instead of 54°C). After each respective PCR program, we used gel electrophoresis to confirm sufficient amplification of each sample and identify any potential contamination using the negative controls. Following PCR2, we prepared samples for Illumina Sequencing by performing a third PCR program that tagged each sample with a unique combination of forward and reverse primers; PCR3 program specifications follow that described above, with the exception of a higher annealing temperature of 60°C. We then normalized the resulting PCR3 product using a SequalPrep Normalization kit (Invitrogen, Burlington, ON, Canada), and shipped the normalized libraries on dry ice for Illumina Sequencing (Illumina MiSeq PE250) at Genome Quebec. Each library was pair-end sequenced in its own lane. On average, each ITS2 library generated 11,970,673 (± 2,893,265) reads, and each rbcL library generated 10,449,713 (± 1,366,310) reads.

**Table 4:**
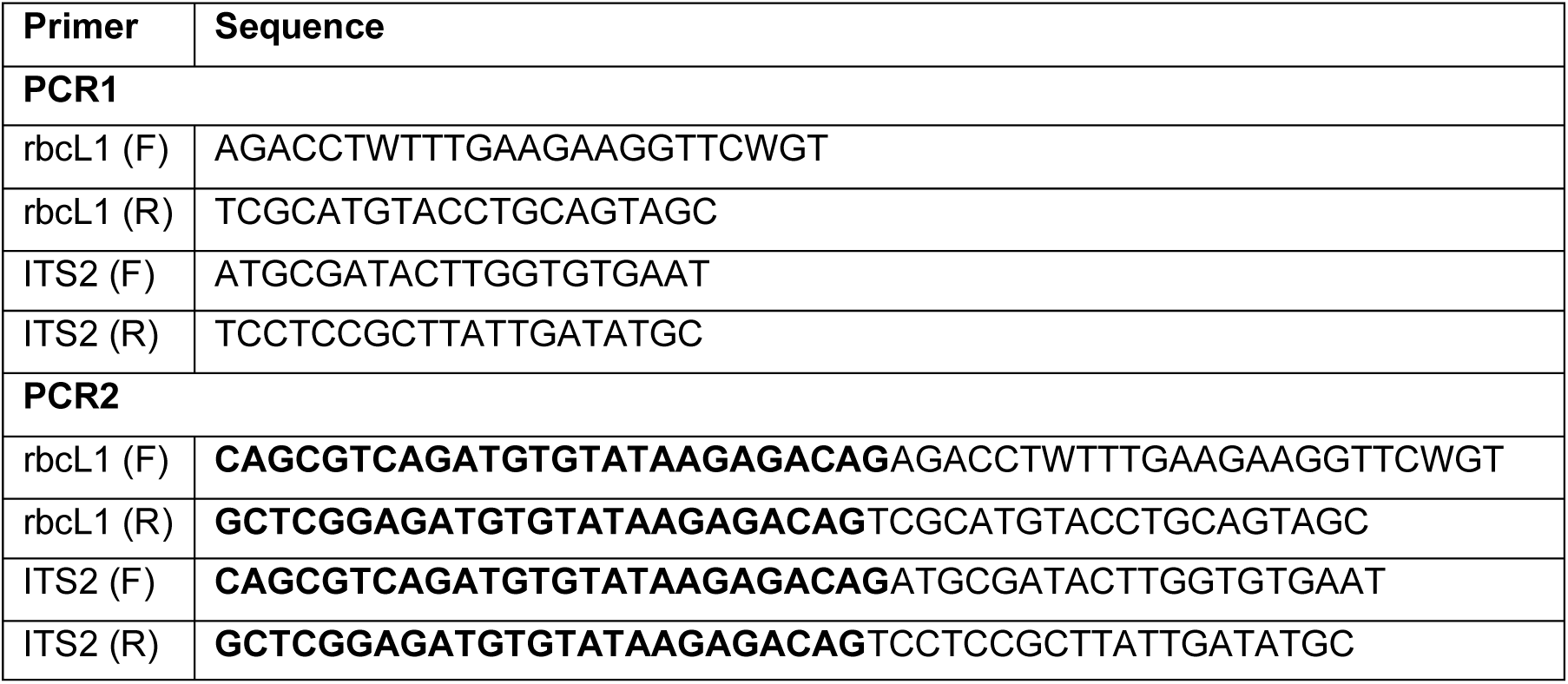
PCR primer specifications. (F) indicates a forward primer and (R) indicates the relevant reverse primer. Bolded values are changes to each respective primer between PCR1 and PCR2 programs.

Pollen metabarcoding data processing was completed in Python (v. 3.10.7), and R (v. 4.2.1; R Core Team), using the *dada2* (83) (v. 1.16.0) and *purrr* (84) (v. 0.3.4) packages. We first paired forward and reverse reads, trimmed primer sequences, and grouped identical sequences as ASVs (amplicon sequence variants). We then built a database that linked species to sequences associated with each primer using the MetaCurator method (85). We used this database to parse through returned sequence data and identify the species associated with each unique grouped sequence, setting a precursory condition of >0.95 similarity. After identifying the plant species associated with each sequence, we consolidated classifications at the genera level, and filtered data to control for sequence mistagging (86). Sequence patterns within negative control samples were used to remove detections with a high likelihood of representing mistag-associated false detections (86).

#### Landscape composition

We used georeferenced raster data from Agriculture and Agri-Food Canada’s 2020 and 2021 Annual Crop Inventory to determine the composition of land cover around sites. The Annual Crop Inventory delineates land cover that was present during the crop growing season for a region each year. Using QGIS (version 3.26.0), we converted the raster data to vector data with polygon features, clipped the land cover data to a 1500-m buffer radius around each apiary location, and calculated the planimetric area of each land cover type. The resulting land type classifications were grouped based on their description. Urban landcover was characterized as the combined total of urban and developed land types. Grassland landcover was a singular group characterized by the grassland landcover type. Forest landcover was characterized as the combined total of coniferous, broadleaf, and mixed-wood land types. Agricultural landcover was characterized as the combined total of all land classifications related to agriculture, including: vegetables, blueberry, cranberry, berry, corn, potatoes, fallow, vineyards, orchards, crops, barley, rapeseed, flaxseed, oats, soybeans, spring wheat, winter wheat, beans, peas, buckwheat, hemp, rye, sugar beets, faba beans, lentils, mustard, triticale, sunflower, millet, and ginseng.

#### Data analysis

All statistical analyses were completed in R (v. 4.2.2, 2022-10-31, R Core Team). To control for sampling effort (i.e. differences in the number of metagenetic sequence hits per sample and its effect on the number of species detected), we filtered each sample to the lowest sequence hit observation for each primer using the ‘rrarefy’ function included in the *vegan* package (87)(v. 2.6-4). After filtering the data, we converted observations into relative frequencies and consolidated the two primers’ estimates to generate multi-locus averages, which are more robust than using single locus estimates (38). We used histograms and residual plots to evaluate the parametricity of response variables and selected subsequent tests based on the distribution. For the focal crop experiments, we used the genus of interests’ multi-locus average relative pollen abundance as our response variable and performed non-parametric repeated measures analysis using a Friedman rank sum test (for within subjects effects across time) and a Kruskal-Wallis rank sum test (for between subjects effects). We treated time as the repeated measure and experimental group (‘near’ vs. ‘far’ crop exposure) as a fixed effect, and when the interaction was statistically significant (p < 0.05), we performed post hoc analysis using Dunn’s test (88) (*dunnstest* package, v. 1.3.5). We calculated Shannon’s diversity indices using the *vegan* package, which were used as the dietary variable for subsequent analyses. For the landscape analysis we used linear models (each site was unique, with little or no experiment-level clustering of landscape parameters), and for stressor analysis we used mixed-effect models. Many of the stressor response variables had insufficient observations (< 25 detections) for either quantitative or logistic modelling, and thus were excluded from analysis. All response variables that met that minimum requirement underwent logistic modelling (10 xenobiotics, 8 pests and pathogens), and a smaller subset that featured both sufficient detection observations and a substantial range of values (e.g. high and low observations) underwent additional quantitative modelling (*Nosema* spores, mites, total summed viral load, thiamethoxam LD_50_ RQ, clothianidin LD_50_ RQ). To account for the influence of time and location, we used experiment (e.g. location x time) as a random effect in all of our models and analyzed the relationship between variables of interest via mixed-effect linear regressions. For our logistic regressions, we used the ‘glmer’ function from the *lme4* package (89) (v. 1.1-31), and for our quasi-poisson regressions, we used the ‘glmmPQL’ function included in the *mass* package (90) (v. 7.3-58.3). We plotted results figures using the *ggplot2* package (91) (v. 4.2.0) with the *ggridges* extension (v. 0.5.4).

#### Data availability statement

All data underlying the findings has been provided as a supplementary file (S2). All code used for analysis has been reposited on GitHub (sbwizenberg/BeeCSI---PNAS-Nexus). We do not provide exact GPS coordinates for apiaries to protect the identity of the beekeepers and farmers involved. The approximate location (municipality, province) of each site is provided in a supplemental file (S2) and further site descriptions are available upon reasonable request.

## Acknowledgments

This work was supported by the Ontario Genomics Institute (OGI-185), Genome Canada and the Ontario Research Fund (LSARP #16420). Supported was also provided by Agriculture and Agri-Food Canada through the Genomics Research and Development Initiative Project (Project J-002368) to SFP, EMW, and MMG. All data underlying the findings is provided as a supplementary file (S2).

