## Supplementary material for "Pollen foraging mediates exposure to dichotomous stressor syndromes in honey bees": S1

**Figure S1**: Shannons Weaver's index of species diversity accurately characterizes dietary composition.

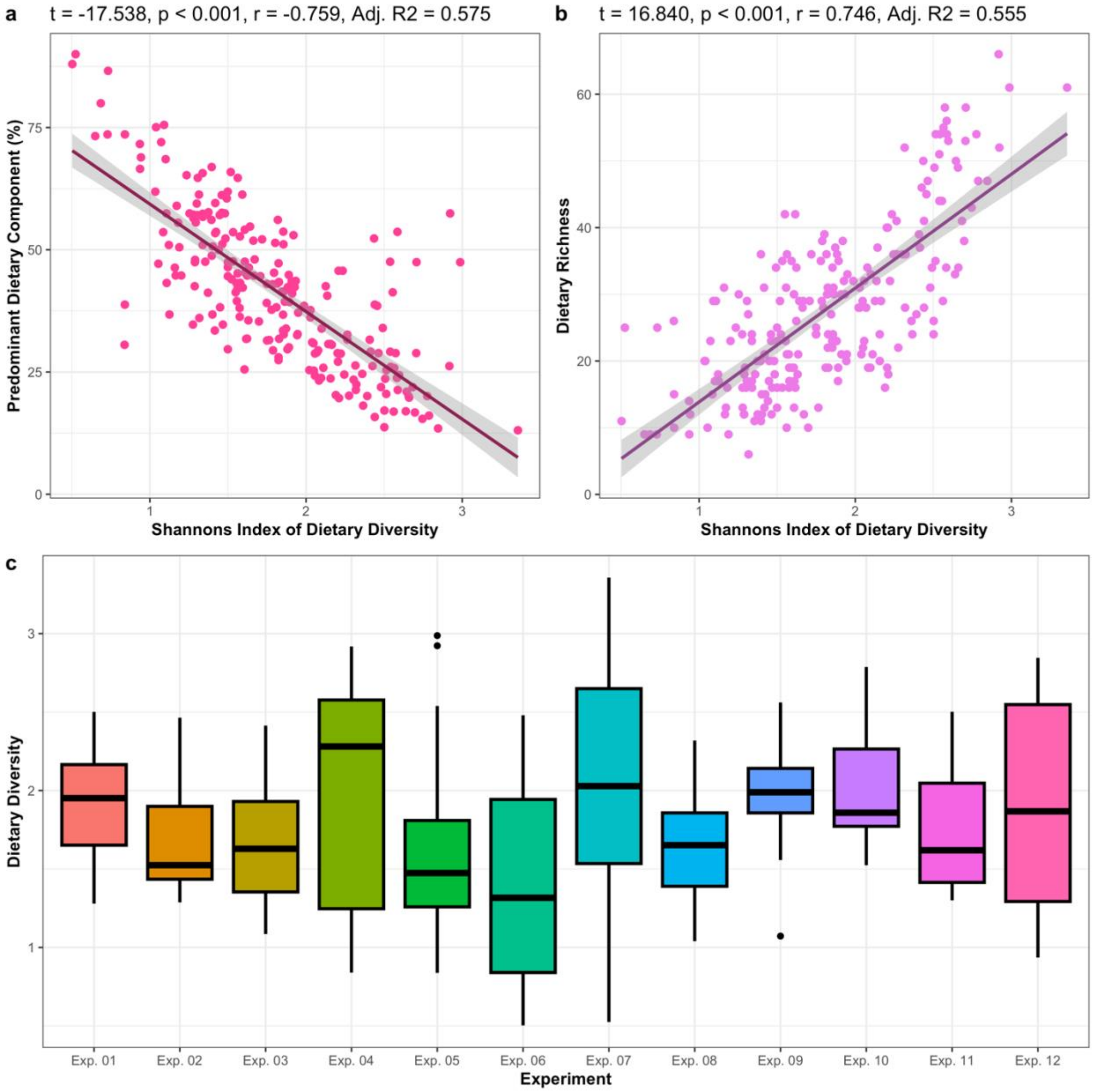


Shannon's index of species diversity, when applied to pollen metagenetic data, (a) correctly indicated the proportion of a diet comprised of the most abundant species (PDC), and (b) correctly indicated the number of species present within each mixed pollen sample. Dietary diversity values varied considerably within experiments (c), providing ideal conditions for mixed-effect general linear modelling.

**Table S1:** Analysis of focal crop exposure experiments using Friedman's repeated measures rank sum, and kruskal-wallis rank sum.

| Response: | | | Focal crop pollen abundance | |
|---|---|---|---|---|
| **Experiment** | **Fixed effect(s)** | **df** | **$X^2$** | **p-value** |
| Apple, 2021 | Time | 1 | 0.000 | 1.000 |
| | Exposure | 1 | 0.145 | 0.703 |
| | Interaction | 3 | 0.296 | 0.961 |
| Corn, 2020 | Time | 1 | - | - |
| | Exposure | 1 | - | - |
| | Interaction | 3 | - | - |
| Soy, 2020 | Time | 1 | 2.000 | 0.157 |
| | Exposure | 1 | 2.105 | 0.147 |
| | Interaction | 3 | 6.316 | 0.097 |
| Canola oil, 2020 | Time | 1 | 1.000 | 0.317 |
| | Exposure | 1 | 0.032 | 0.859 |
| | Interaction | 3 | 0.474 | 0.925 |
| Canola oil, 2021 | Time | 1 | 6.400 | 0.011 |
| | Exposure | 1 | 0.052 | 0.820 |
| | Interaction | 3 | 5.049 | 0.168 |
| Canola seed, 2020 | Time | 1 | 10.000 | 0.002 |
| | Exposure | 1 | 0.023 | 0.879 |
| | Interaction | 3 | 14.589 | 0.002 |
| Canola seed, 2021 | Time | 1 | 3.600 | 0.058 |
| | Exposure | 1 | 1.120 | 0.289 |
| | Interaction | 3 | 5.377 | 0.146 |
| Cranberry, 2020 | Time | 1 | 0.500 | 0.479 |
| | Exposure | 1 | 3.944 | 0.047 |
| | Interaction | 3 | 14.358 | 0.002 |
| Cranberry, 2021 | Time | 1 | 0.143 | 0.706 |
| | Exposure | 1 | 7.011 | 0.008 |
| | Interaction | 3 | 9.401 | 0.024 |
| Highbush blueberry, 2020 | Time | 1 | 0.667 | 0.414 |
| | Exposure | 1 | 2.004 | 0.157 |
| | Interaction | 3 | 4.819 | 0.186 |
| Highbush blueberry, 2021 | Time | 1 | 2.000 | 0.157 |
| | Exposure | 1 | 3.081 | 0.079 |
| | Interaction | 3 | 11.059 | 0.011 |
| Lowbush blueberry, 2021 | Time | 1 | 5.000 | 0.025 |
| | Exposure | 1 | 6.169 | 0.013 |
| | Interaction | 3 | 18.506 | < 0.001 |

$X^2$ is the test statistic, and p-value indicates the statistical significance of the relationship ($\alpha = 0.05$). Significant p-values are bolded.

**Table S2**: Post hoc analysis of focal crop exposure experiments using Dunn's rank-sums test of multiple comparisons with a benjamini-hochberg adjustment.

| Experiment | Pairwise comparison | Second grouping | p-value |
|---|---|---|---|
| Canola seed, 2020 | Exposed vs. unexposed | t1 | 0.395 |
| | | t2 | 0.378 |
| | t1 vs. t2 | Exposed | 0.007 |
| | | Unexposed | 0.016 |
| Cranberry, 2020 | Exposed vs. unexposed | t1 | 0.290 |
| | | t2 | 0.001 |
| | t1 vs. t2 | Exposed | 0.004 |
| | | Unexposed | 0.164 |
| Cranberry, 2021 | Exposed vs. unexposed | t1 | 0.248 |
| | | t2 | 0.010 |
| | t1 vs. t2 | Exposed | 0.134 |
| | | Unexposed | 0.223 |
| Highbush blueberry, 2021 | Exposed vs. unexposed | t1 | 0.448 |
| | | t2 | 0.015 |
| | t1 vs. t2 | Exposed | 0.008 |
| | | Unexposed | 0.560 |
| Lowbush blueberry, 2021 | Exposed vs. unexposed | t1 | 0.500 |
| | | t2 | 0.001 |
| | t1 vs. t2 | Exposed | < 0.001 |
| | | Unexposed | 0.750 |

P-value indicates the statistical significance of the relationship ($\alpha = 0.025$). Significant p-values are bolded.
